# Big data analysis of mitochondrial DNA substitution models: A regression approach elucidating the effects of codon position and neighboring nucleotides

**DOI:** 10.1101/272922

**Authors:** Keren Levinstein Hallak, Shay Tzur, Saharon Rosset

## Abstract

We build on the up-to-date version of Phylotree, a comprehensive and continuously updating phylogeny of global human mtDNA variations (van Oven and Kayser 2009), to better understand the substitution mechanism of the mitochondrial DNA (mtDNA) and its most influential factors. We do so by composing Poisson and negative-binomial regression models relating the rate of occurrence of mtDNA substitutions to various factors. Important factors we identify include the identity of the codon at each position, confirming previous findings about the biological significance of different codons for the same amino acid. Importantly, we also identify a significant effect of neighboring sites. This effect cannot be attributed solely to CpG pairs. A similar effect of neighboring sites was recently described for autosomal DNA substitutions, and we speculate it is related to the basic mutational mechanism itself. Once codon composition and context are taken into account, there is no significant difference in substitution rate between different genes in mtDNA.

## Introduction

The human mitochondrial DNA (mtDNA) is a short circular haploid and non-recombinant chromosome, which is maternally inherited. It also has different codons, replication and proof reading mechanisms compared to the autosomal DNA (Anderson et al. 1981; Clayton 2000; Johnson and Johnson 2001). The lack of recombination allows for a simpler analysis of the complex substitution process underlying the phylogenetic mechanism, compared to recombinant DNA. Specifically, all historical human mtDNA can be modeled using a common phylogenetic tree, with the leaves specifying extant individuals. Branchings in the tree describe separations of lineages that lead to different offspring today. The root of the tree can be designated as “mitochondrial Eve” (Cann, Stoneking, and Wilson 1987). The process of inferring the tree structure from sampled DNA sequences has been studied extensively in the literature where methods such as maximum likelihood (Felsenstein 1981), maximum parsimony (Czelusniak et al. 1990) and neighbor joining (Saitou and Nei 1987) were proposed and compared (Kolaczkowski and Thornton 2004; Takahashi and Nei 2000). Several software packages containing these methods are publicly available (Koichiro Tamura et al. 2011; Yang 1997).

There are a few comprehensive mtDNA databases such as MITOMAP (Kogelnik et al. 1996), mtDB (Ingman and Gyllensten 2006) and Phylotree (van Oven and Kayser 2009) that are regularly updated. In this work we use the highly trusted phylogenetic tree reconstructed by Phylotree. Phylotree has been updated several times throughout the years and currently (Build 17) consists of 24,275 sequences (reference files) and 5,437 nodes. Behar et al. (2012) published a refinement of the tree which included a high confidence reconstruction of its root which they termed RSRS. In our work we use the most updated tree and the RSRS root, and assume it trustfully describes the true phylogenetic history. Since most of the observed mtDNA sequences are located in the leaves, we can refer explicitly to three elements of potential uncertainty in generating the list of substitutions inferred from the tree: First, we assume that the phylogeny of the tree specifying its topological structure as given by Phylotree is correct. This assumption was previously justified in several papers (Behar et al. 2012; Röck et al. 2013; Rosset et al. 2008; Soares et al. 2009) and is mainly supported by the sparsity of substitutions in branches in deeper layers of the tree and the resulting successful haplogroup estimations.

The second source of uncertainty relates to the ancestral sequences in the hidden layers of the tree. Most nodes can be confidently reconstructed by maximum parsimony, however there may still be some uncertainty in the reconstruction (Rosset et al. 2008).

Finally, considering the identity and number of specific substitutions along the branches, it is possible that several substitutions occurred in one site on the same branch, but the sequences in the nodes only contain the initial and final state of the site on this branch. For example, in a certain site there might be a C base transitioning to T, followed by a transversion to A. It may be the case that only the starting C and finishing A are estimated in the nodes’ sequences, thus underestimating the number of substitutions in that site by assuming only one *C*→*A* transversion occurred. A variety of similar events can happen, where multiple substitutions follow in a branch in the same site. Soares et al. (2009) argue that in a long branch in the human mtDNA phylogeny, the probability that one of the highly mutated sites in mtDNA (site 152) has multiple substitutions on the branch is below 1%. Since most of our analyses do not consider such fast sites at all, the probability of repeated unobservable substitutions in our analysis is probably negligible. Subsequently, we feel confident in assuming all substitutions are specified by Phylotree. This greatly simplifies the statistical analysis as it requires no additional inference on latent information.

In this work we focus on genic regions in mtDNA, though our methods can be easily extended to other sub-regions. We use the recent increase in data to perform comprehensive statistical tests that were previously impossible. Our data driven approach focuses on analyzing the significance of the effect of possible factors on the substitution rate. Relevant factors might include the current base/codon, the sub-region in which the site is located, its neighboring nucleotides and more. Assessing the importance of each factor has a crucial effect on understanding the underlying biological principles affecting mtDNA substitutions. For example, if some sub-regions are likely to behave according to the same substitution model, then from a biological view they are likely to be functionally related and from a statistical standpoint we can aggregate these sub-regions to increase the power of future research.

### Our main contributions are as follows

1. We utilize the most updated Phylotree data and execute an exhaustive analysis on the effect of multiple factors on the substitution count process through both Poisson and negative-binomial regressions. We examine for each factor if it should be included as an explaining variable in the substitution count process and if the substitution model should be partitioned according to it. The factors we examine include categorical factors that are not constant per site (for example: the input codon, amino acid, nucleotide and context vary in each site). We incorporate these factors into the regression by adding for each factor an exposure term that takes into account the time spent in each site for each possible value of the categorical factor. We also examine if it is beneficial to model all substitutions together or separate them into transitions/transversions or synonymous/non-synonymous substitutions. To the best of our knowledge this is a new statistical perspective in the context of DNA data allowing to quantify the significance of all factors simultaneously. A detailed description of our approach is given in the Methods section.
2. Our results show that the substitution model should be partitioned into sub-models according to the codon position and input codon; each sub-model should include the neighboring sites and direction of replication as explaining variables. Genes, however, should not be included in the model at all. In addition, we show it is advantageous to model transitions and transversions together as well as synonymous and non-synonymous substitutions.
3. We apply a novel clustering technique on genes that is based on five similarity tests between each pair of genes as detailed in the Methods section. Our new method supports previously found gene functionalities.

### Previous Works and Points of Interest

There is a plethora of works that deal with substitution models for DNA. Extensive literature consider a reversible continuous Markov chain model such as JC69 (Jukes and Cantor 1969), F81 (Felsenstein 1981) and most notably HKY85 (Hasegawa, Kishino, and Yano 1985). These models describe the probability for every base to transition to every other base after a specific time duration using a rate matrix whose constraints differ between the possible models. When considering only the number of substitutions per site, models with substitution rates that are independent of the current nucleotide (such as JC69 and F81) induce a Poisson distribution on each site.

Subsequently, more recent works study site dependent rates. When the rate of each site is sampled from a Gamma distribution, the marginal distribution can be described using a negative binomial (NB) distribution as was done by Tamura and Nei (1993). Hence, the most natural way to embed additional factors into the substitution probabilities is to redefine the rate per site as a variable dependent on the relevant factors, for example via a Poisson regression. In case the substitution rate is not determined by the considered factors, but rather has an additional unexplained variability (that might stem from additional factors we have not considered), the substitution rate can be modeled using a NB regression. Indeed, in our work we assume the rate of each site can be modeled using either a Poisson or a negative-binomial regression, where achieving a good model via Poisson regression would be considered generally better as it implies we have considered all relevant factors.

A different body of works deals with more specific testing of different factors and phenomena in DNA substitution models such as hot-spots (Galtier et al. 2006), CpG pairs (Lunter and Hein 2004) and context (Aggarwala and Voight 2016; Siepel and Haussler 2004). For instance, context is considered an important factor in coding sequence non-randomness utilized for efficiency and accuracy in protein synthesis (Fedorov 2002). In their work, Aggarwala and Voight (2016) showed that most of the variability in polymorphism levels in autosomal DNA can be attributed to the context, and used these results to detect irregularities correlated with neurodevelopmental and psychiatric disorders. Indeed our results also show that context is a significant explaining variable, even when other explaining variables are considered. Most similar to our work is the work of Zoller and Schneider (2010), who empirically investigated what are the most relevant parameters for a codon model by applying principal component analysis (PCA) on more than 3,000 codon substitution rate matrices estimated from single gene families. Their results showed that the first principal component (PC) separates synonymous and non-synonymous substitutions while the second PC distinguishes between substitutions where only one nucleotide changes and substitutions with two or three nucleotide changes. Our approach goes beyond theirs in that it is parametric and allows direct interpretation of the different factors’ effects and significance; we also concentrate on mtDNA.

Some works consider the codon structure instead of relating directly to bases. Several codon-substitution models based on the reversible continuous Markov chain model have been suggested, where usually the key parameter is the damping of non-synonymous transition rates (Yang and Nielsen 2008). In the regression model we test, we consider both the codon and the amino acid it encodes as relevant factors and test their effect on the substitution rate. Subsequently, in some sense we test whether the basic unit of the model should be a base, a codon or an amino acid, while enveloping all options as viable. There is a vast literature comparing nucleotide, amino-acid and codon based models (Kosiol and Goldman 2011; Seo and Kishino 2009; Simmons 2017; Whelan et al. 2015), all works but the last support codon based models. Indeed, our results agree that codons are the basic unit required for inference on the number of substitutions.

Site-based partitioning of the substitution model into independent substitution sub-models unfolds the assumption that each group of sites has evolved under the same evolutionary process (independently of the other groups). Partitioning was applied ad hoc in several works, mainly according to codon position and genes (Ho and Lanfear 2010; Nylander et al. 2004; Shapiro, Rambaut, and Drummond 2006). Due to the intractable nature of performing an exhaustive search over all partitioning options, several partitioning search algorithms were developed, such as a hierarchical clustering method suggested by Li et al. (2008), and the popular PartitionFinder open source program (Lanfear et al. 2012) which efficiently finds optimal partitions using a heuristic search algorithm. In this work we chose to implement an exhaustive search over all possible partitioning options for several reasons; first, heuristic searches cannot be guaranteed to find the optimal partitioning scheme (Li et al. 2008) and thus if computationally possible, an exhaustive search is always preferred. Second, the previously mentioned methods perform site-based partitions, allowing to partition over factors such as genes and codon position. However, factors that change along the tree in each site such as nucleotide, amino acid, codon and neighboring nucleotides cannot be incorporated into this site-based partitioning method.

## Results

### Genes Clustering

In the Methods section we suggest several similarity tests to examine the null hypothesis that two genes are similar in terms of sequence statistics (tests 1-2) and dynamical behavior statistics (tests 3-5). These tests were applied to each pair of genes and their p-value was compared to a Bonferroni corrected critical value of 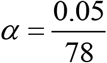. Our results are summarized in Table 1. Under all tests, the null hypothesis could not be rejected for pairwise comparison of genes from the group: *ND1-5*, *CO2* and *CO3* suggesting that they should be clustered together. When considering only dynamical behavior tests (tests 3-5) this group can be expanded to contain *ND1-6* and *CO1-3*. From here on, we shall refer to this group of genes (*ND1-6* and *CO1-3*) as *NDCO*. *ATP6* and *ATP8* can be joined under all tests and will be referred to as *ATP*.

For the purpose of modeling the substitution process of mtDNA we have decided to use groups formatted by the dynamical statistics tests only, resulting in the following groups: *NDCO*, *ATP* and *CYB*. We note that our clustering procedure was done without taking into consideration the biological functionality underlying the different sub-regions (Anderson et al. 1981), subsequently, the tests we performed further support the previously found partitions. The sequence statistics tests (tests 1-2) show that ND6 is different from other ND genes. This cannot be explained by the different directionality of ND6, as after changing ND6 sequence content to its Watson-Crick complement (Watson and Crick 1953) we still reject the null hypothesis when comparing ND6 to other genes in NDCO according to tests 1-2.

**Table 1.** The results of the clustering tests on the different pairs of sub-regions; each cell in the table contains the indices of the null hypotheses which were rejected (ranging from 1 to 5). Tests 1/2 test the null hypothesis that the distributions of the nucleotides/codons are the same in RSRS, whereas tests 3-5 compare the substitution count distribution through (3) Kruskal-Wallis, (4) negative-binomial model and (5) negative-binomial regression. Tests 1 and 2 are not directly related to the dynamics of the model, so the resulting clusters by our tests are (*ND1-6, CO1-3*), (*ATP6, ATP8*) and *CYB*.

|  | ND1 | ND2 | ND3 | ND4L | ND4 | ND5 | ND6 | CO1 | CO2 | CO3 | ATP6 | ATP8 | CYB |
| --- | --- | --- | --- | --- | --- | --- | --- | --- | --- | --- | --- | --- | --- |
| ND1 |  |  |  |  |  |  | 1 2 |  |  |  | 3 4 | 3 4 |  |
| ND2 |  |  |  |  |  |  | 2 | 1 2 |  |  | 3 4 5 | 3 4 | 3 4 |
| ND3 |  |  |  |  |  |  | 1 2 |  |  |  | 3 4 5 | 3 4 |  |
| ND4L |  |  |  |  |  |  | 2 |  |  |  | 3 4 5 | 3 4 | 3 4 5 |
| ND4 |  |  |  |  |  |  | 2 | 1 2 |  |  | 3 4 5 | 3 4 | 3 4 5 |
| ND5 |  |  |  |  | NDCO |  | 1 2 | 1 2 |  |  | 3 4 5 | 3 4 | 3 4 |
| ND6 |  |  |  |  |  |  |  | 1 2 | 1 2 | 1 2 | 2 3 4 | 3 4 | 1 2 |
| CO1 |  |  |  |  |  |  |  |  |  |  | 3 4 5 | 1 3 4 | 3 4 5 |
| CO2 |  |  |  |  |  |  |  |  |  |  | 3 4 5 | 1 3 4 | 3 |
| CO3 |  |  |  |  |  |  |  |  |  |  | 3 4 5 | 1 3 4 |  |
| ATP6 |  |  |  |  |  |  |  |  |  |  | ATP |  | 3 4 |
| ATP8 |  |  |  |  |  |  |  |  |  |  |  |  |  |
| CYB |  |  |  |  |  |  |  |  |  |  |  |  | CYB |

### Exhaustive Search Algorithm

We aim to find the best substitution model, where we use Akaike information criterion (AIC) (Akaike 1974) as a criterion for comparing models. We use the term model to denote a specific choice of inclusion for each one of the categorical variables that might affect the substitution rate as listed in the Methods section. The inclusion options for each variable are as follows: for each categorical explaining variable we can choose between (1) not including it in the model, (2) including it as an explaining variable or (3) partitioning the data according to it and building a separate sub-model for each part of the data. Each model is therefore composed of a varying number of sub-models according to its specific inclusion options. For example, a model that partitions the data according to the codon position and does not include all other variables will be composed of three sub-models — one for each codon position.

We calculated the AIC for each one of these models with the output as (1) the sum of all substitutions seen in each scenario, (2) two separate models with combined likelihood, one for the number of synonymous substitutions and the other for the number of non-synonymous substitutions observed at each scenario and (3) two separate models with combined likelihood, one for the number of transitions and the other for the number of transversions observed at each scenario.

Overall we examined 31,185 models and ordered them by their minimal AIC score (obtained by either Poisson regression, or NB regression); the top 20 results of our algorithm are given in Table 2 and all results appear in Supplementary Table S2. For each model we specify which factors partitioned the model into sub-models (marked as 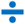), and which factors were included / not included in all the resulting sub-models (marked as 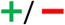 correspondingly).

All top 20 models:

- Partition into sub-models according to the codon position
- Include the input codon (as an explaining variable or partition according to it) and subsequently do not include the input amino acid and input nucleotide
- Include the direction of replication (as an explaining variable or by partition)
- Do not divide the type of substitution into separate models for transitions/transversions or synonymous/nonsynonymous substitutions.
- Result in a substantially lower AIC for the NB model compared to the Poisson model

Also, most of the top 20 models include the right and left neighbors.

The model with the lowest AIC score is composed of 192 sub-models with different codon positions and input codons (64 sub-models for each of the three codon positions, hence 192=64·3 sub-models). The explaining variables in each model are the right and left nucleotide neighbors, the directionality and the additional site-based variables that were included in all models.

Another interesting result relates to the subject of model partitioning which was discussed in the introduction. As previously mentioned, there are several algorithms that heuristically find best fit partition schemes (Lanfear et al. 2012; Li et al. 2008). While these algorithms necessarily result in site-based partitions, our results clearly show that the model with the lowest AIC score partitions the data according to the input codon in addition to the codon position. Indeed, previous works (Ho and Lanfear 2010; Shapiro, Rambaut, and Drummond 2006) have shown that partitioning the substitution model according to the codon position is beneficial. However, partitioning according to the input codon with additional conditions was not considered yet due to practical limitations (Kalyaanamoorthy et al. 2017), though it is shown here to be advantageous.

We examined the effect of neighboring nucleotides and CpG pairs as explaining variables and found that neighboring nucleotides have a significant effect and should be included as explaining variables in the model. Our definition of neighboring nucleotides refers to nucleotides outside of the codon when the input codon is included in the model, so for the first codon position we take into account only the left neighbor, for the second codon position no neighbors are considered and for the third codon position only the right neighbor is considered.

Comparing models one and five in Table 2 (that differ only by the inclusion of the neighboring nucleotides) using a generalized likelihood ratio (GLR) test allows to examine the *H*_0_ hypothesis that the neighboring nucleotides have an insignificant effect on the substitution rate. The result (p-value <1*e*^−12^) shows that the neighboring nucleotides have a significant effect on the substitution rate. However comparing models five and six (that differ only by including the CpG trait as an explaining variable) using a GLR test shows that the CpG trait effect alone is insignificant. Note that the leading model partitions the data according to the input codon and codon position, and also includes neighbors, so it technically contains the CpG trait information. To conclude, our results show that neighboring nucleotides have a significant effect on the substitution rate and should be added as an explaining variable, whereas including the CpG trait alone is not enough.

### Poisson Vs. Negative Binomial Regression

The leading models show AIC scores that are substantially lower for NB regression than for Poisson regression. As stated before, NB regression is preferable when there are missing relevant factors and a latent Gamma-like distribution replaces the uncertainty, resulting in a more appropriate NB regression; so this indicates that there might be additional factors we have not considered that affect the substitution rate. Such factors can be for example the clade in which the substitution occurred, the time if the molecular clock assumption is not valid or perhaps an inherent randomness suggesting that a NB model (or perhaps another model that takes this randomness in substitution rate into account) is indeed preferable compared to a Poisson model.

It is important to note that even if the leading model had a lower AIC score for the Poisson model than the NB model, it would still not mean necessarily that we have found all relevant factors. To see why, consider for example model #258 as ranked by the AIC score (appears in Supplementary Table S2): this model has the lowest Poisson AIC score out of all models and its NB AIC score is significantly higher than its Poisson AIC score (*AIC*=30,049 for the Poisson model and *AIC*=48,021 for the NB model). The model partitions the data according to codon positions, directionality and genes and includes the CpG pair trait and input codon as explaining variables. If we had not considered the first 257 models, this would have been our leading model suggesting that we have considered all relevant factors. However this is clearly not the case as our leading model has a lower AIC score, and its AIC score is lower for the NB regression.

**Table 2.**
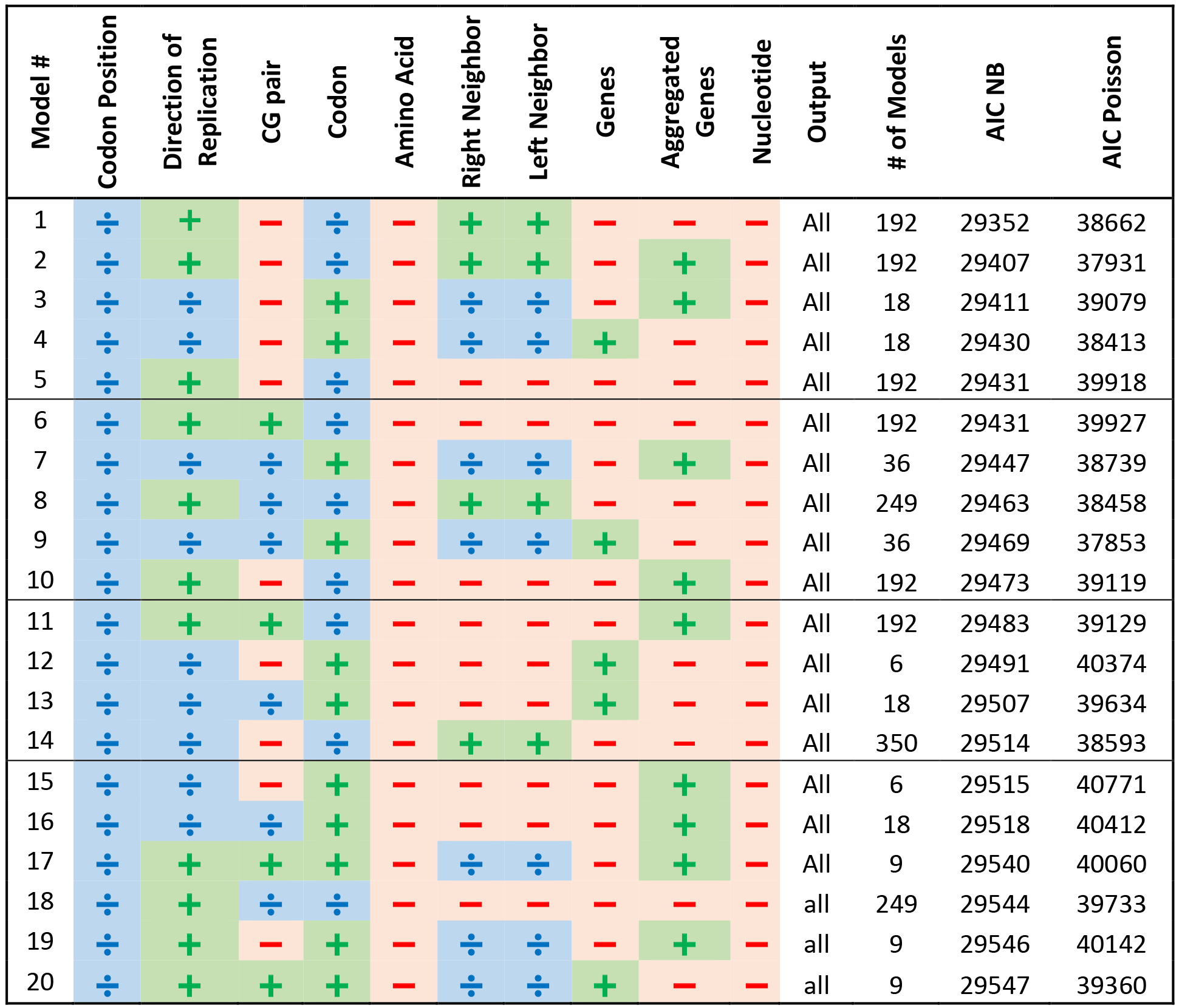
First 20 models ordered by their minimal AIC score (out of Poisson and NB AIC scores). Each categorical variable obtains one of the following signs: 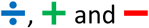 that mark partitioning, inclusion as an explaining variable and omitting the variable correspondingly. The value all in the output column means all substitutions were modeled together (and not separately for transitions/transversions and synonymous/non-synonymous substitutions.

## Discussion

Despite the extensive research in the field, there is limited understanding of the factors affecting substitution rates in various DNA modalities, and specifically in mtDNA. In our view, the reason for that is relatively small amounts of data, compared with the possibly large number of degrees of freedom stemming from the various possible factors affecting the substitution rate: site location, haplogroup association, non-stationarity, codon properties, context and latent biochemical information. In this work we utilized the recently updated Phylotree to tackle this problem using proper statistical tools.

Unlike continuous time Markov chain models which specify the transition rate between every two instantiations of the basic model units (nucleotides, codons or amino-acids), we model directly the distribution of the substitutions count in a time interval for a specific set of observed factors. This formulation allows for simple Poisson/NB regressions to simultaneously consider combinations of variables participating as either partitions or as explaining variables. We can thus choose the most suitable model through the likelihood and degrees of freedom for each suggested model.

The model with the lowest AIC score includes partitions according to the input codon and the position inside the codon. Neither the amino acid itself, nor the input nucleotide, were enough to hold all information required to model the substitution process. Particularly, the neighboring sites have a significant effect in setting the substitution process rate coefficients. We note that the observed context significance refers to neighboring sites adjacent to the codon (and not inside the codon). This significance remains even when the CpG trait is considered as an explaining variable, so the CpG trait alone is not sufficient to explain the context effect. Subsequently, in future works an expanded context should be considered as well. While the effect of nucleotide substitutions in amino-acid codons can be explained by natural selection constrains as a result of protein structure and function, the origin of the detected effect described herein of neighboring sites cannot be explained by protein level constrains and we speculate it is related to the nature of mutagenesis in mtDNA. For example, an imperfect replication process by mtDNA polymerase γ (POLG) that is related to the neighboring sites, and suggested to be responsible for the majority of mtDNA point mutations (Szczepanowska and Trifunovic 2017). Nonetheless, the specific biological mechanism causing this effect is yet to be identified.

As far as we are aware, the direction of replication, relevant only for ND6, was never considered explicitly as an explaining variable. This could be due to the fact that genes are commonly considered as explaining variables and thus include the direction of replication as an explaining variable as well. Our results show its effect can be captured sufficiently in an additive fashion, simplifying the model.

Interestingly, the genes themselves were not significant enough, which somewhat discourages the notion of purifying selection. Were the substitution rate affected by the functionality of the gene itself, then probably the more crucial genes would have shown a reduced substitution rate.

We plan to extend this work to the other regions of the mtDNA, while adding the haplogroup and time as additional explaining factors, where our main goal is finding a unified substitution model which can improve our understanding of the underlying biological mechanisms.

The statistical approach we present here can be further applied to autosomal DNA, with appropriate adjustments considering its different properties; the recombination process in autosomal DNA challenges the possibility of finding a clear phylogeny allowing to track substitutions over time. However, its substitution rate is much slower compared to the mtDNA and is almost unique-event polymorphism (UEP). Approximating the substitution rate to be UEP, we can use logistic regression to model the probability that a substitution occurred over time, allowing to examine which factors are significant in the autosomal substitution process.

## Methods

### Data Preprocessing

The substitutions data was obtained from the Phylotree website (van Oven and Kayser 2009), where the tree was constructed so that only substitutions that were shared by at least three complete sequences were included (with a few exceptions). The substitutions A16182c, A16183c, C16519T/T16519C were not considered for phylogenetic reconstruction and these sites are therefore excluded from the data. In the phylogeny tree there are 12,961 substitutions and 5,437 leaves — mtDNA sequences that represent a haplotype found in human populations. There are 1,113 leaves that have one representative mtDNA sequence (reference file) and 3,578 that have two. We examined these sequences and found 20,696 substitutions that were not included in the tree (most of them singletons) and added them to the data, resulting in a total of 33,657 substitutions. There are overall 16,056 substitutions in genic regions which are the focus of this work. The empirically observed transition/transversion ratio is 21.97 in the mtDNA overall and 22.4 in the genic regions. The empirically observed nonsynonymous/synonymous substitution ratio is 0.418.

The approach taken in this paper requires filtering the substitutions by additional parameters other than the site in which they occurred. For example, we are interested in filtering substitutions according to the neighboring nucleotides. These filtrations are achievable by first forming for every node in the tree the full mtDNA sequence it represents, and then for every substitution in each branch store the corresponding parameters.

### Genes Clustering

Some genes are known to have similar functionality, which they were found and named according to. Aggregation of genes that follow the same substitution model effectively reduces the number of distinct sub-models required to characterize the data, thus reducing the degrees of freedom and increasing the power of performed statistical tests. Yet, blindly aggregating genes together by functionality can lead to aggregating genes with different substitution models, or to missing a previously unknown similarity in substitution models between different genes.

To better support any choice of aggregation, we have devised several similarity tests between genes. Each test examines the hypothesis that two genes share some mutational or sequence characteristics and returns an appropriate p-value. The combined results for all tests can either support or oppose the aggregation of every two genes. The tests are explained as follows:

1. Compare the distributions of nucleotides in the genes sampled on the RSRS sequence, using a two-sample *χ*^2^-test.
2. Compare the distributions of codons in the genes sampled on the RSRS sequence, using a two-sample *χ*^2^-test.
3. Compare the per-site substitution count of the two genes using Kruskal-Wallis test (Kruskal and Wallis 1952).
4. Fit a negative-binomial model to the number of substitutions per site for each gene separately, and for the aggregation of the two genes. Compare the results using a GLR test.
5. Perform negative-binomial regression with the gene as an explaining variable — its p-value corresponds to whether the genes should be joined.

The first two tests above compare the sequence statistics of the two genes while the rest test the dynamic behavior under different varying harshness of assumptions. Note that tests four and five are very much alike and test similar null hypotheses but they are not identical. If several genes indeed behave identically with respect to the substitution model, we expect none of the hypotheses to be rejected between every pair of these genes. We applied each one of the five above mentioned tests to each pair of sub-regions 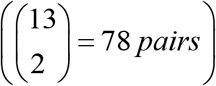 and compared the obtained p-values to a Bonferroni corrected critical value 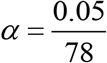. A p-value lower than this threshold means that the null hypothesis that the two compared genes are “the same” (the meaning of this is different for each test) was rejected. We used the results to cluster together genes that their comparison yielded p-values higher than the critical value. The rule we applied for clustering genes was that in order to be joined, two genes must be “the same” under all tests that refer to the substitution model (i.e. tests 3-5). The resulting clustered genes groups are as follows (see Results for details):

1. ***NDCO****: ND1, ND2, ND3, ND4L, ND4, ND5, ND6, CO1, CO2, CO3*
2. ***ATP****: ATP6, ATP8*
3. ***CYB***

### Variables affecting the substitution rate

The models consider the following categorical factors:

1. Genes
2. Clustered genes as described in the previous section
3. The input nucleotide which was present before the substitution occurred (A/C/G/T)
4. The input amino-acid which was present before the substitution occurred (21 amino-acids)
5. The input codon which was present before the substitution occurred (64 codons)
6. The codon position (1/2/3)
7. The right and left neighboring nucleotides
8. Whether or not the site was part of a CpG pair and if so was it the first or second position
9. Directionality (indicator to whether or not the gene is located on the light strand; considering mtDNA protein coding genes its value is 1 only for ND6)

Additional site-based factors included in all models are evolutionary conservation as calculated by phyloP100way vertebrate (Pollard et al. 2010) and protein family, transmembrane and low complexity domains as found by Bateman et al. (2002).

### Poisson and Negative Binomial regressions

Poisson point process is a memoryless count process implying exponential waiting time between events. Assuming that the substitution process is a Poisson process requires the assumption that it is memoryless, i.e. that the number of substitutions at each time interval is independent of the number of substitutions in non-overlapping time intervals. The substitution rate, which we will denote here as *λ*, may however depend on both observed and unobserved variables. Examples for possibly relevant observed variables are gene, codon position and other variables mentioned in the previous subsection. When all relevant variables are observed and the memoryless assumption holds, the substitution process can be modeled as a conditional Poisson process. If we also assume that the dependence of log(*λ*) on the variables is linear, then log(*λ*) can be approximated as a linear combination of the input explanatory variables by applying a Poisson regression with a log link function. The optimal solution can be found through Fisher’s scoring method (Longford 1987) or otherwise by convex optimization schemes (such as gradient descent). It might be the case that in addition to the relevant observed variables, there are also latent variables (variables that were not deemed as relevant, or hidden variables that were not observed at all) affecting the substitution rate. In this case the substitution process can be properly modeled as a conditional over-dispersed Poisson process.

If the over-dispersion is properly modeled by a gamma function, the resulting substitution model follows a NB distribution. The NB model loosens the Poisson assumption of equal mean and variance, by introducing an unobserved heterogeneity term for each site. The notion of underlying Gamma distributed rates for every site was previously considered by Tamura and Nei (1993) and became a standard approach in inference over the phylogenetic tree (Yang 1994; Yang 1997; Koichiro Tamura et al. 2011).

We follow this view and perform both Poisson regressions and NB regressions to model the substitution rate at each site. If the Poisson model is less preferable (we use AIC to score models), then this means there are still missing relevant factors and a latent Gamma-like distribution describes the uncertainty, resulting in a better fitting NB regression.

### Adding exposure to the Poisson and Negative Binomial regressions

We define a state *S* as a specific set of values of the explaining variables discussed above. For example, the state *S*_*Site*=7765,*Codon*=*GAG*, *Neighbor* = *A*_ corresponds to site 7765 with an input codon GAG and right neighbor A. Note that once the site, codon and neighbor are known, all other variables are also known (the site determines the relevant gene, clustered gene, codon position, directionality and the additional site-based factors; the codon determines the amino acid and when the codon position and neighbor are known the input nucleotide and CpG condition can also be determined).

Assuming the substitution process is a Poisson or NB process dictates that the expected number of substitutions is proportional to the amount of time spent in each state. We therefore calculated the time spent in the tree at each state and added this time as an exposure variable; for instance, in order to include the input codon as an explaining variable, we calculated for each site how much time was spent in each codon over the whole tree. While this could potentially augment the data times 64, in fact at each site there were no more than four different codons throughout the tree. Adding the time as an exposure variable to a regression with a log link function (such as the Poisson and NB regressions we applied) means that the regression finds the optimal *β* coefficients such that log(*λ*) = *β*^*T*^ *x* + log(*time*), forcing *λ*, the expected number of substitutions to be proportional to the time spent at the state as required.

We also add an exposure variable to the case where we separate synonymous and non-synonymous substitutions. When the number of synonymous substitutions is modeled separately, there are states in which a synonymous substitution is impossible. Consider for example the codon GTT that codes for the Valine (Val) amino acid: in its first and second position there are no possible synonymous substitutions, while in its third position there are three possible synonymous substitutions — one transition (GTT → GTC) and two transversions (GTT → GTA, GTG). In the same manner we can calculate the number of possible non-synonymous substitutions for each codon position and divide them into possible transitions and transversions. We expect the number of synonymous substitutions at each state to be proportional to the number of **possible** synonymous substitutions with respect to the number of possible transitions and transversions. Subsequently, when the number of synonymous/non-synonymous substitutions was modeled separately we added the number of possible synonymous/non-synonymous substitutions as an exposure explaining variable, when taking into account the number of possible transitions/transversions and weighting them according to the transversions/transitions ratio empirically found in the data (*r* = 0.04551103). In these cases the regressions for synonymous substitutions find the optimal *β* coefficients such that: log(*λ*) = *β*^*T*^ + log(*time*) + log(#*A* + *r* ·#*B*), where #*A* is the number of possible synonymous transitions and #*B* is the number of possible synonymous transversions.

### Time Estimation

In order to find the exposure of each factor’s value, we require an estimate of the time-length of each branch in the tree (or alternatively, the time of each branching event in the tree). As proposed by (Rosset 2006), we used a Poisson regression with an identity link function for time inference on all nodes in the tree. Substitutions that occurred in the tips of the tree were assigned the time *t*=0. Notice that the assigned times are uncalibrated — since the exact timing of any branching in the tree is unknown, only the proportions between timings can be inferred. This also implies that all obtained substitution rate estimates are relative — all of them can be multiplied by a constant and all time estimates divided by the same constant without changing the likelihood. Table S1 in the supplementary material contains uncalibrated time estimations for each node in the Phylotree dataset.

### Exhaustive Search Algorithm

We examined 31,185 models composed of three different inclusion options for each categorical variable. The number of examined models is not three to the power of the number of factors since there are many exceptions where models are removed when they are contained in another model. For once, we did not consider models with both codon and amino acid factors, nor models with both codon and input nucleotide factors, since the codon variable contains the information given by both; hence the number of possible models resulting from variation of the codon, amino acid and input nucleotide inclusion options is 11 instead of 27. For the same reason we did not consider models with both the gene and clustered gene so the number of possible inclusion options for the gene and clustered genes variables is five instead of nine. When the data was partitioned according to the codon position the right and left neighbor variables were coerced to follow the same choice of inclusion. Hence, the number of possible inclusion options for the codon position and right and left neighboring nucleotides is 21 instead of 27. We also note that for models with the codon as an explaining variable which were partitioned according to the codon position we included only the right and left neighboring nucleotides outside of the codon (left neighbor for the first codon position, right neighbor for the third codon position and none for the second codon position). Finally, some of the resulting models were removed from the analysis since the inclusion of certain factors contained all information on the value of another factor (for example genes contain information about the directionality since only ND6 has opposite directionality). Such models were removed from Table 2.

Overall we examined 11·5·21·3^3^ = 31,185 models, calculated as the product of number of options for each variable as specified before: codon / amino acid / input nucleotide, gene / clustered gene, codon position / neighboring nucleotides, directionality, CpG pair status and the possible output models.

We note that this exhaustive search over all options was necessary since the inclusion of each categorical variable may affect the other variables. For example, if the model is partitioned into sub-models according to the input codon, it may no longer be statistically useful to divide into sub-models according to the gene. We finish this section with some technical remarks: The 31,185 models we examined are composed of 8,460,470 sub-models when considering the partitions according to all categorical variables; so we performed 8,460,470 NB and Poisson regressions, out of which 51,291 (0.006%) NB regressions and 18,062 (0.002%) Poisson regressions did not converge. Since all sub-models are needed for calculating the AIC score of each model, it was necessary to include the models that did not converge. In order to do so, we assigned these models likelihood according to a Poisson distribution with a parameter *λ* equal to the number of observed substitutions in each state and the degrees of freedom were taken to be the number of different states (rows) in the sub-model. In addition there are 3,212,855 sub-models that are composed of one state or include zero substitutions, whose likelihood and degrees of freedom were taken to be 1 (except for sub-models that include one state with at least 1 substitution whose likelihood was calculated according to a Poisson distribution).

## Acknowledgments

This work was supported in part by a fellowship from the Edmond J. Safra Center for Bioinformatics at Tel-Aviv University.

